# Spermidine and spermine are the natural substrates of the *Acinetobacter baumannii* AmvA multidrug efflux pump

**DOI:** 10.1101/2020.10.02.324624

**Authors:** Francesca L. Short, Qi Liu, Heather E. Ashwood, Varsha Naidu, Liping Li, Bridget C. Mabbutt, Karl A. Hassan, Ian T. Paulsen

## Abstract

Multidrug efflux pumps are important drivers of antibiotic resistance in *Acinetobacter baumannii* and other pathogens, however their ‘natural’ roles beyond transport of clinical antimicrobials are poorly described. Polyamines are an ancient class of molecules with broad roles in all three kingdoms of life, and are the likely natural substrate of at least one efflux pump family. We have defined the transcriptome of *A. baumannii* following treatment with high levels of the polyamines putrescine, cadaverine, spermidine and spermine. These molecules influenced expression of multiple gene classes in *A. baumannii* including those associated with virulence, and the four polyamines induced distinct but overlapping transcriptional responses. Polyamine shock also induced expression of the MFS-family efflux pump gene *amvA* and its repressor gene *amvR*. Loss of *amvA* dramatically reduced tolerance to the long-chain triaamine spermidine, but caused only modest changes in resistance to known AmvA substrates such as acriflavine. We confirmed reduced accumulation of spermidine in *amvA*-deficient *A. baumannii*, and showed that its expression is induced by long-chain polyamines through its cognate regulator AmvR. Our findings suggest that the conserved *A. baumannii* efflux pump AmvA has evolved to export spermidine from the cell, but that its substrate recognition promiscuity also allows activity against clinically-important biocides and antibiotics.

**Importance:** AMR genes, including multidrug efflux pumps, evolved long before the ubiquitous use of antimicrobials in medicine and infection control. Multidrug efflux pumps often transport metabolites, signals and host-derived molecules in addition to antibiotics or biocides. Understanding the ancestral physiological roles of multidrug efflux pumps could help to inform the development of strategies to subvert their activity. In this study, we investigated the response of *Acinetobacter baumannii* to polyamines, a widespread, abundant class of amino acid-derived metabolites, which led us to identify long-chain polyamines as natural substrates of the disinfectant efflux pump AmvA. A second clinically-important efflux pump, AdeABC, also contributed to polyamine tolerance. Our results suggest that the disinfectant resistance capability that allows *A. baumannii* to survive in hospitals may have evolutionary origins in the transport of polyamine metabolites.

## Introduction

Antimicrobial resistance is a critical public health challenge of the 21^st^ century. Infections caused by antibiotic-resistant bacteria are predicted to cause 10 million annual deaths by 2050 if urgent action is not taken (1). Though extensive AMR is a clinical phenomenon, the genes responsible have deep evolutionary origins long predating the use of antibiotics in medicine (2). Naturally-occurring antibiotics are typically found at subinhibitory levels, while synthetic drugs and biocides are not present in natural (unpolluted) microbial environments at all. As such, AMR genes are proposed to play additional roles, for example in bacterial signalling, metabolism and virulence (3, 4). Multidrug efflux pumps are an important category of AMR determinant, and often have core physiological functions (5, 6). These functions broadly fall into removal of harmful exogenous molecules (for example, mammalian antimicrobial peptides or plant flavonoids), or secretion of endogenous molecules (such as siderophores, quorum-sensing signals or metabolites). Understanding the origins and ancestral functions of AMR genes, including MDEs, could help to predict their future evolution, or point to new ways to subvert their activity.

*Acinetobacter baumannii* is a notorious opportunistic pathogen classified within the “ESKAPE” group of bacterial species that are responsible for the majority of antibiotic-resistant infections (7, 8). *A. baumannii* has a particular ability to survive for prolonged times in hospital environments, due in part to high disinfectant tolerance conferred by its repertoire of multidrug efflux pumps (9, 10). Many *A. baumannii* efflux pump genes are conserved across the species or genus, suggesting a shared primordial function. Recently it was shown that the physiological substrates of AceI – a chlorhexidine efflux pump encoded on the core genome of *A. baumannii* ^19,20^ – are likely to be short-chain polyamines (11).

Polyamines are an ancient class of metabolites comprising two or more amine moieties connected by an aliphatic chain; the most common biological polyamines being putrescine, cadaverine, spermidine and spermine (Fig 1A)(12). These molecules have central roles in all three kingdoms of life, and can be present intracellularly at high (mM) concentrations (12, 13). In bacteria, polyamines have been implicated in species-specific functions which include biofilm formation, cell growth, oxidative stress resistance, and nitrogen storage, among others(14). Many pathogenic bacteria depend on polyamine synthesis or import for their pathogenesis (15–18). Polyamines can exert their functions in bacteria directly by virtue of their general biochemical properties, or they can serve as signals which act through specific receptors even at low concentrations (17, 19). Whether produced endogenously or present in the environment, polyamines can be toxic in high amounts. Several efflux systems have been reported to facilitate polyamine transport, such as members of the small multidrug resistance (SMR)(20), major facilitator superfamily (MFS)(21), and PACE families (11).

**Figure 1.**
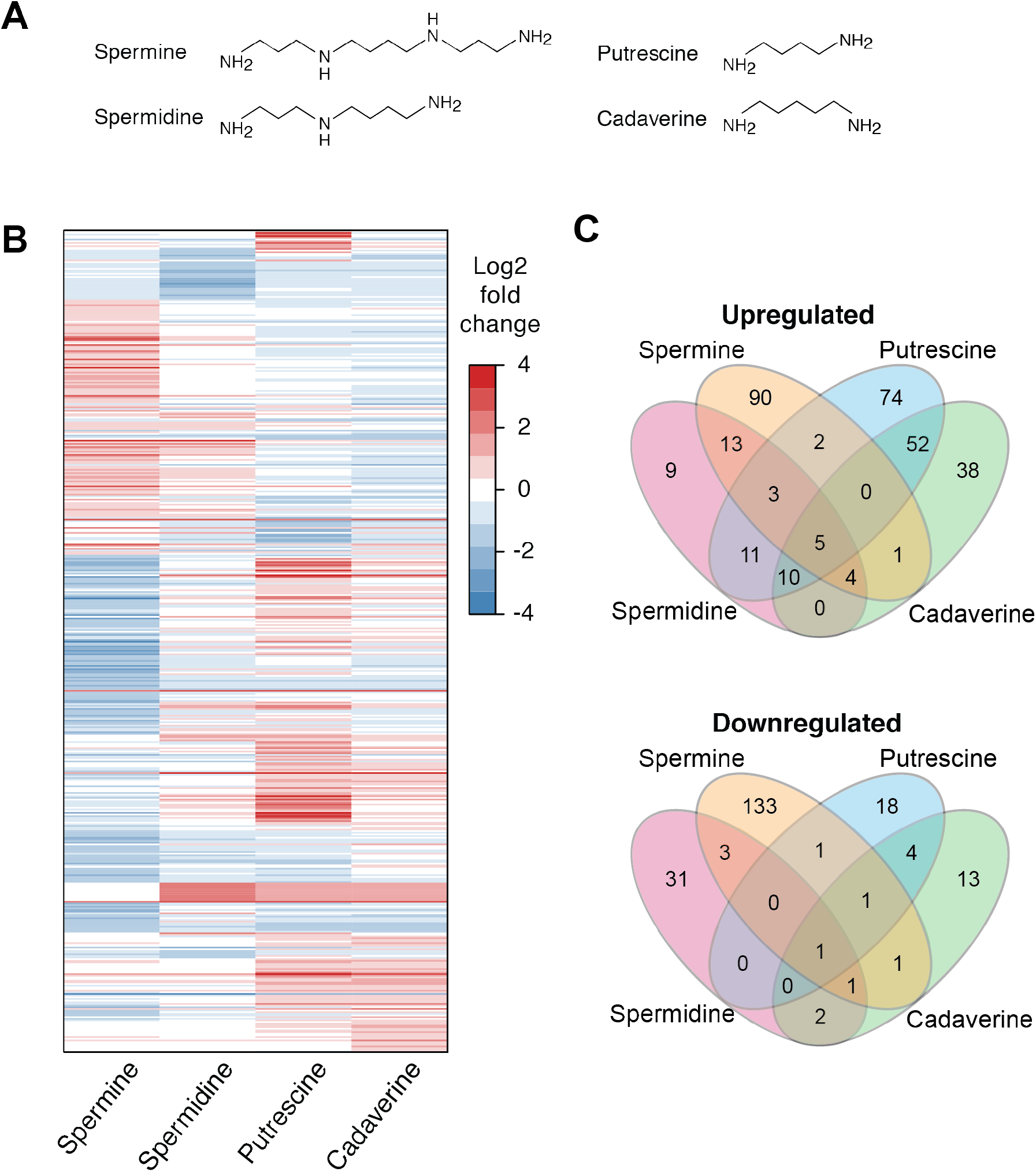
Distinct regulons of spermine, spermidine, putrescine and cadaverine. (A) Structures of the four polyamines used in this study. (B) Heatmap of expression changes following polyamine shock, showing the direction of expression change, and concordance between different polyamines. (C) Venn diagram of up- and downregulated gene sets. Note that some targets showed expression changes in opposite directions following treatment with different polyamines.

In this study, we have investigated the transcriptomic response of *A. baumannii* to high levels of the four major biological polyamines (putrescine, cadaverine, spermidine and spermine), with a view to defining their physiological roles, and identifying transporters responsible for efflux of these molecules. The efflux pump genes *aceI*, *adeABC* and *amvA* were all strongly induced by different polyamines. We show that AdeABC and AmvA are required for tolerance to the long-chain polyamines spermine and spermidine, and demonstrate transport of spermidine by the MFS transporter AmvA. Finally, we also show that spermine and spermidine induce *amvA* expression by direct binding of its repressor, AmvR. Our results strongly suggest that long chain polyamines are physiological substrates of the conserved *A. baumannii* efflux pump AmvA.

## Results

### RNA-Seq of A. baumannii following polyamine shock

We performed transcriptomics on *A. baumannii* AB5075-UW (Table S1) following shock with exogenous polyamines. The minimum inhibitory concentrations of all molecules were very high at 10 mg/ml for spermine and 40 mg/ml for spermidine, putrescine and cadaverine. RNA was extracted from duplicate log-phase *A. baumannii* AB5075-UW cultures supplemented with putrescine di-hydrochloride, cadaverine di-hydrochloride, spermidine tri-hydrochloride, or spermine tetra-hydrochloride at 1/8 MIC. RNA-Seq reads were mapped, normalised and fold-changes calculated as described (see Materials and Methods). Each biological replicate gave rise to >10 million reads with >98% mapping to the *A. baumannii* AB5075-UW genome (Table S2). Genes showing significantly altered expression in the presence of polyamines were defined as those with log2 fold change >1 or <-1 and a corrected p-value <0.05 relative to the control.

### Individual polyamines induce distinct transcriptional responses

A total of 499 genes showed altered expression following treatment with one or more polyamines, with individual regulons ranging from 93 (spermidine) to 259 (spermine) genes (Fig 1B; Table S3). The diamines putrescine and cadaverine each caused upregulation of a large number of genes (157 and 110, respectively) and downregulation of relatively few (25 and 23). For the tetraamine spermine and triamine spermidine, the number of genes showing increased and decreased expression was more balanced (spermine: 118 up/141 down; spermidine: 55 up/38 down). Though the majority of gene expression changes were specific to just one polyamine (372 genes), there was substantial overlap in the genes regulated by putrescine and cadaverine, which had 73 common targets (Fig 1B, C). Interestingly, putrescine and spermine showed divergent regulation of 21 genes. These genes included ABUW_0068-71 (involved in amino acid metabolism), ABUW_2096-2099 (fatty acid metabolism) and ABUW_2448-2456 (fatty acid metabolism). Nine genes were differentially expressed with all four polyamines: the *adeABC* efflux pump genes, the transcriptional regulator *amvR* (ABUW*_*1678; also called *smvR*), and ABUW*_*0233 were induced, the periplasmic OB-fold protein-encoding ABUW_1352 was repressed, and three genes (ABUW_2448, ABUW_2449 and ABUW-2453) were induced by putrescine, cadaverine and spermidine but repressed by spermine.

### Functional categories of polyamine-responsive genes

A summary of the COG functional classifications of polyamine-responsive genes is shown in Fig 2. Enrichment of specific GO terms within the polyamine-regulated genes was tested using the TopGO package (22). A summary of polyamine-regulated genes of interest is given in Table S4. Frequent targets included metabolic genes, particularly those involved in energy production and conversion, lipid transport and metabolism, inorganic ion transport, and amino acid metabolism (COGs C, I, P and E, respectively).

**Figure 2.**
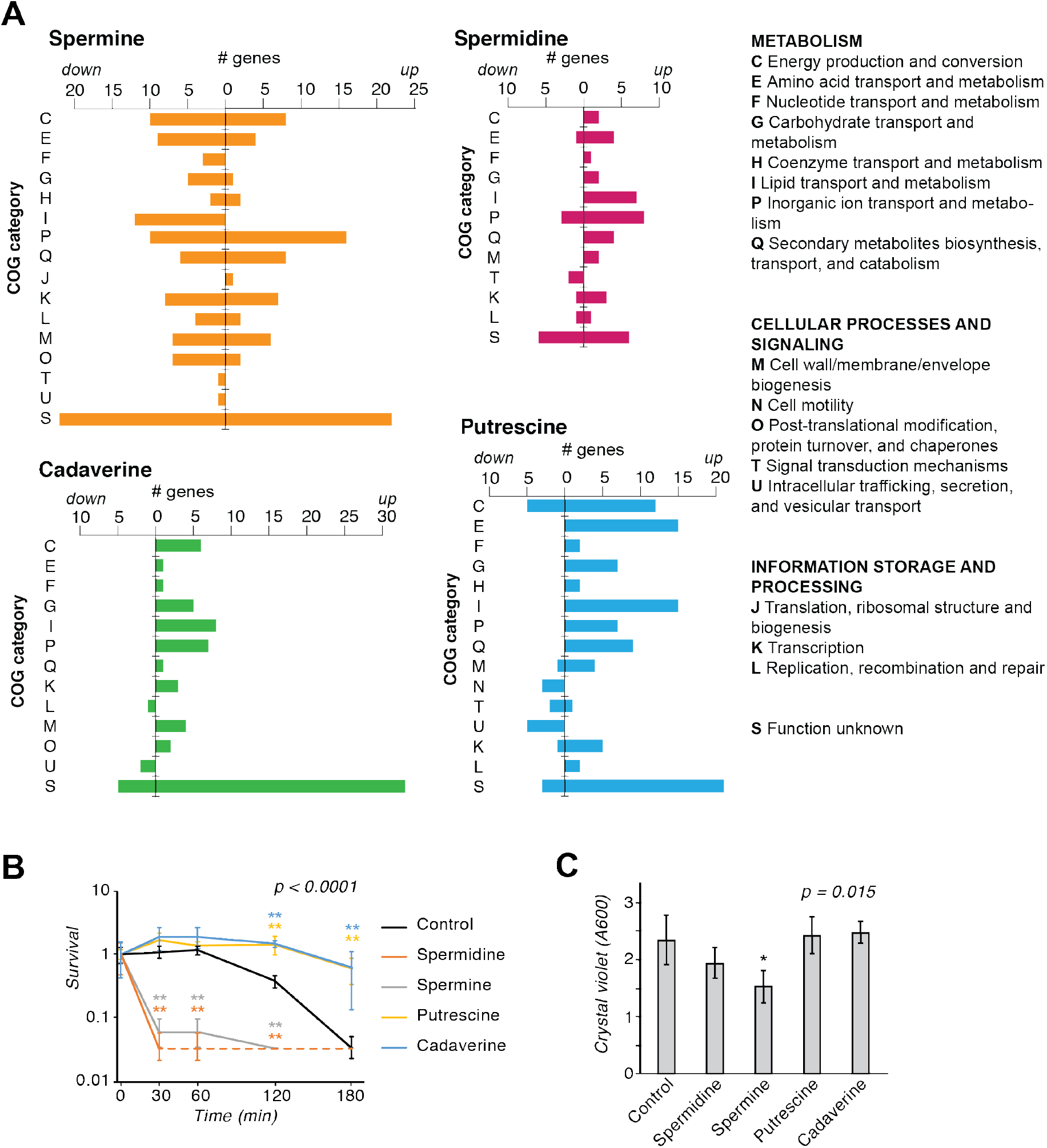
Polyamine-regulated functional categories and phenotypes. (A) Functional categories of genes regulated by spermine, spermidine, putrescine and cadaverine. COG categories of the genes showing expression changes in response to each molecule are shown. Regulated genes came from multiple functional classes, and lipid/inorganic ion transport and metabolism genes were highly represented. (B) Serum survival curves following treatment of 106 cells with 66− normal human serum supplemented with polyamines at 1/8 MIC (n = 3). Statistical significance was determined by two-factor repeated measures ANOVA on log10-transformed survival data (overall p-value < 0.0001) followed by a one-factor ANOVA with Dunnett’s post-hoc test at individual time points to compare all samples to the control. **p < 0.01. (C) Biofilm formation in the presence of polyamines (n = 3). Static 2.5ml cultures with or without polyamine supplementation were incubated for 40 hours at 37C, and the attached biomass quantified by crystal violet staining. *p < 0.05, one-way ANOVA with Dunnett’s post-hoc test (overall p-value = 0.015).

Transcription genes (COG K) were also common polyamine-responsive targets, particularly with spermine.

As expected given that polyamines are amidated molecules and their metabolism is tightly linked to that of amino acids, several operons involved in utilisation of amino acids were regulated by polyamine shock. This included genes for histidine utilisation (induced by putrescine and spermidine), leucine metabolism genes ABUW_2453-5 (induced by putrescine and cadaverine, repressed by spermine), and many amino acid transporters. Polyamine shock also regulated genes involved in fatty acid utilisation such as genes of the *ato* butyrate/acetoacetate degradation operon (induced by putrescine and cadaverine), ABUW_1572-4 (induced by spermidine, putrescine and cadaverine), and the ABUW_2447-58 operon for leucine and fatty acid degradation.

### Regulation of virulence genes and phenotypes in response to polyamine shock

Virulence of *A. baumannii* depends on multiple factors including secretion systems, pili, siderophores, efflux pumps and other defense systems (23).

Multiple genes from virulence-related pathways were induced or repressed by exogenous polyamines (Table S4). Many iron-acquisition genes were polyamine-responsive, such as the siderophore synthesis and uptake genes induced by spermidine and spermine. In addition, polyamines downregulated the expression of several virulence-related cell surface structures including the *csu* pilus (suppressed by cadaverine), genes of the Type VI secretion system used for interbacterial competition (repressed by cadaverine and spermidine), and Type IV secretion system and competence genes (suppressed by putrescine). Polyamine-regulated stress resistance genes included those for copper resistance (induced by spermine, putrescine and cadaverine), heavy metal resistance (induced by cadaverine and spermine), and hydrogen peroxide resistance. Finally, polyamines appeared to affect expression of genes involved in horizontal gene transfer, as well as horizontally-acquired elements themselves.

In addition to the Type IV pili mentioned above, which are required for horizontal gene transfer in *A. baumannii* (23), some enzymes of the CRISPR-Cas phage resistance locus were repressed by spermine, and expression of genes within predicted prophage regions (24) was variably affected by spermidine and putrescine (Table S4).

We conducted serum resistance and biofilm formation assays to determine whether the transcriptional effects determined above would drive changes in virulence-related phenotypes. *A. baumannii* AB5075-UW showed a delayed susceptibility to serum killing, with a loss of viability between 1 and 3 hours (Fig 2B). Spermidine and spermine at 1/8 MIC both caused a drastic reduction in serum survival that was apparent after 30 minutes. Unexpectedly, putrescine and cadaverine supplementation increased serum survival to close to 100−. *A. baumannii* AB5075-UW showed relatively low biofilm formation after 40-hour static incubation in rich medium (Fig 2C). The presence of putrescine, cadaverine or spermidine at 1/8 MIC did not affect static biofilm formation in rich media, while spermine caused a small decrease. These results do not support a major role for polyamines in biofilm regulation in *A. baumannii* AB5075-UW despite transcriptional regulation of some relevant genes (eg. the *csu* pilus). While the opposing effects of short-chain and long-chain polyamines on serum resistance are consistent with their divergent transcriptional regulation, we did not notice any specific regulatory targets that would explain their effects on serum resistance.

### Multidrug efflux pumps regulated by polyamine shock

*A. baumannii* encodes a wide repertoire of multidrug efflux pumps, which confer the ability to withstand many disinfectants and antibiotics. Drug efflux pumps are often induced by their substrates (5). The known efflux pump genes *aceI*, *adeABC* and *amvA* all showed increased expression in the presence of polyamines, with different combinations inducing each one (Fig 3A). The PACE family transporter gene *aceI* was the most dramatically upregulated, with a log2-fold change of ~8 in the presence of putrescine and cadaverine, and 2-4 with spermidine and spermine. This fits with the recent finding that short chain diamines (including putrescine and cadaverine) are physiological substrates of AceI, while the longer-chain polyamines spermine and spermidine are not (11). The *adeABC* genes, encoding an RND family transporter, showed a 2-3 log2-fold increase in expression in the presence of any of the four molecules. The MFS transporter gene *amvA* was induced by spermidine and spermine but not by putrescine and cadaverine. The *amvA* repressor gene *amvR* (25) was also induced by polyamine shock, however the cognate regulators of *aceI* and *adeABC* (*aceR* and *adeRS*, respectively) were not.

**Figure 3.**
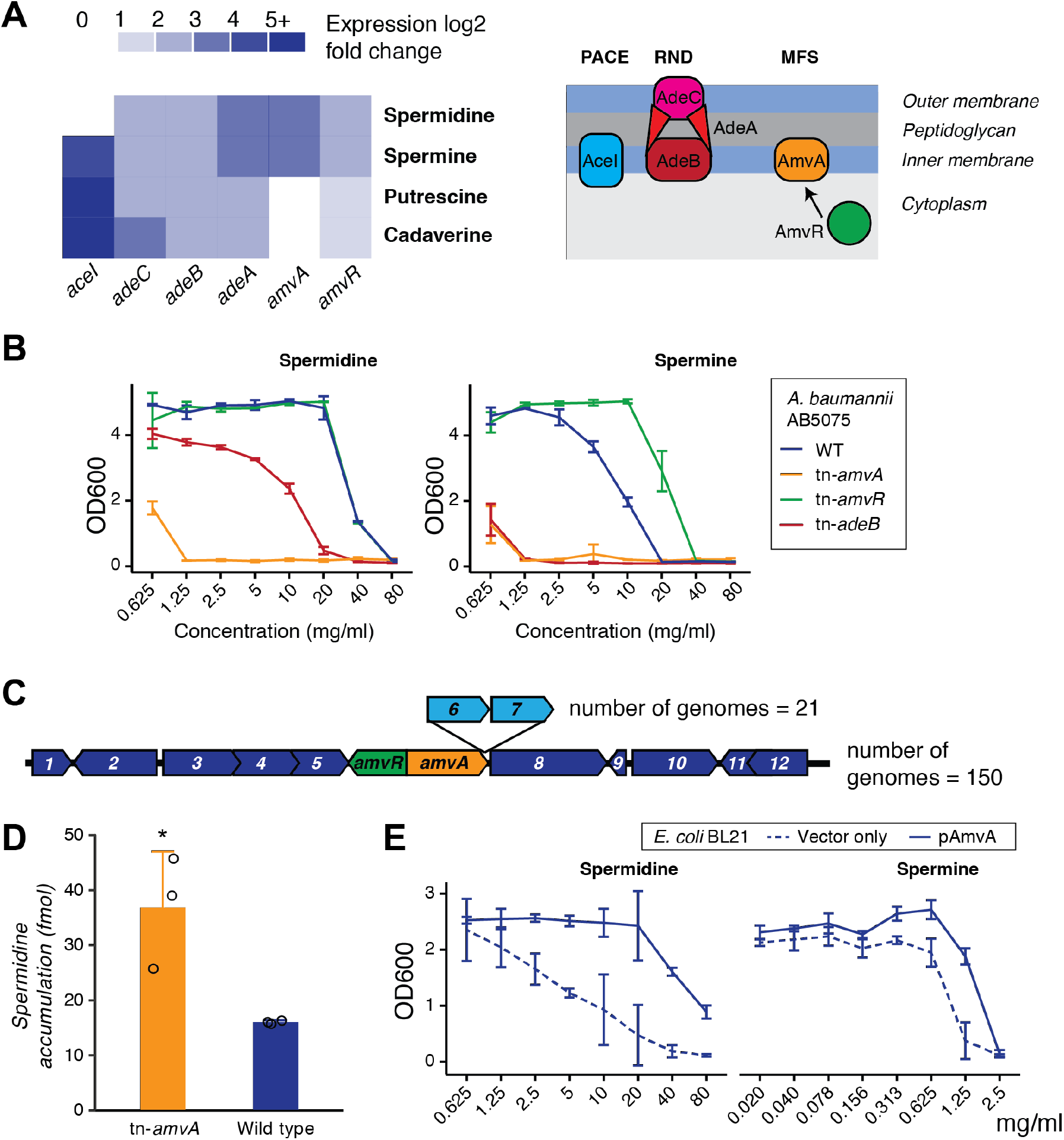
Induction of efflux pumps by polyamine shock identifies spermine and spermidine as AmvA substrates. (A) Discontinuous heatmap showing induction of the efflux pump genes *aceI* (PACE family), *adeABC* (RND family) and *amvA* (MFS family) and its regulator *amvR* by different polyamines, with a schematic of these efflux pumps to the right. (B) Growth of *A. baumannii* AB5075 and mutants in the presence of spermine and spermidine. Results are mean */− SD for two biological and two technical replicates. (C) Genomic context of *amvAR* in 172 *A. baumannii* strains. The genes are present in 100− of complete *A. baumannii* genomes in a region of very high synteny. Core genes are shown in dark blue, accessory genes in light blue, and the number of genomes possessing each gene path is indicated. Annotated gene functions are: 1 – *aceI* PACE family transporter, 2 - acetyl-CoA acyltransferase, 3 - short-chain dehydrogenase/ reductase, 4 - MaoC domain protein dehydratase, 5 - beta-lactamase, 6 - hypothetical protein, 7 - hypothetical protein, 8 - *dnaX* DNA pol III subunit tau, 9 - hypothetical protein, 10 - phospholipase C, 11 - thioesterase family protein, 12 - iron-containing alcohol dehydrogenase. (D) Intracellular accumulation of 3H-spermidine in *A. baumannii* AB5075 and its tn-*amvA* mutant. Late exponential-phase cells were washed into assay buffer and incubated with 10.8 nM 3H-spermidine (see Materials and Methods), and the amount of accumulated radioactivity measured by liquid scintillation counting. Results shown are from three independent biological replicates. * p< 0.05, one-way ANOVA. (E) Growth of *E. coli* BL21 in the presence of spermidine and spermine, with or without AmvA overexpression. Results are mean */− SD for three biological and two technical replicates.

### AmvA is a transporter of long-chain polyamines

As there are no characterised spermine or spermidine efflux pumps in *A. baumannii*, we sought to determine whether AmvA and/or AdeABC may perform these roles. First, growth in the presence of polyamines was measured to test whether AmvA or AdeABC may be responsible for removing harmful levels of these molecules from the cell (Fig 3B; Table 1). Mutation of *amvA* dramatically decreased *A. baumannii* AB5075-UW resistance to spermine (16-fold reduction in MIC) and spermidine (64-fold reduction). Mutation of *amvR*, which increases *amvA* expression (25), increased the MIC of spermine 2-fold to 20 mg/ml. An *adeB* mutant showed an 8-fold reduction in spermine MIC and a 2-fold reduction in spermidine MIC. Tolerance to putrescine and cadaverine was not affected by *amvR*, *amvA* or *adeB* mutation. These observations support the hypothesis that spermine and spermidine may be the natural substrates of the well-characterised drug efflux pumps AmvA and AdeABC.

**Table 1.** Polyamine MIC changes in *amvA*, *amvR* and *adeB* mutants of *A. baumannii* AB5075

| *Polyamines* | **WT** | MIC (mg/ml) [fold-change] |  |  |
| --- | --- | --- | --- | --- |
|  |  | *tn-amvA* | *tn-amvR* | *tn-adeB* |
| Spermine | 10 | 0.625 [16] | 20 [0.5] | 1.25 [8] |
| Spermidine | 40 | 0.625 [64] | 40 | 20 [2] |
| Putrescine | 40 | 40 | 40 | 40 |
| Cadaverine | 40 | 40 | 40 | 40 |

| *Reported AmvA substrates* | **WT** | MIC (ug/ml) [fold-change] |
| --- | --- | --- |
|  |  | *tn-amvA* |
| Acriflavine | 64 | 16 [4] |
| Benzalkonium chloride | 32 | 32 |
| Erythromicin | 250 | 250 |
| Ethidium bromide | 32 | 32 |
| Methyl viologen | 100 | 25 [4] |

AmvA is found in all *A. baumannii* genomes, within a region of very high synteny comprised of other conserved genes (Fig 3C). Due to its very strong phenotype, and its absolute conservation hinting at a deeply-rooted role in *A. baumannii* biology, AmvA was selected for further characterisation. An *amvA* mutant of *A. baumannii* AB5075 was tested against a selection of its previously reported substrates (26) in order to compare its effect on resistance to these molecules with its effect on polyamine resistance (Fig 3B; Table 1). Mutation of *amvA* did not affect the MIC of three of the five substrates tested, while two substrates – acriflavine and methyl viologen – showed a 4-fold MIC reduction in AB5075-UW tn-*amvA*. Note that previous investigation of the AmvA substrate range used *A. baumannii* strain AC0037; AB5075-UW has a different complement of efflux pumps which could potentially mask some AmvA activities. Overall, AmvA has a much greater impact on resistance to toxic levels of polyamines than on resistance to its previously reported substrates.

We then tested whether loss of *amvA* affects accumulation of spermidine in cells (Fig 3D). *A. baumannii* AB5075-UW and *A. baumannii* AB5075-UW *tn-amvA* were grown to exponential phase, washed, and incubated with 10.8 pmol [^3^H]-spermidine. Mutation of *amvA* significantly increased the amount of intracellular [^3^H]-spermidine, though accumulation of exogenous spermidine was very low in the conditions tested (approx 15 fmol in 2 × 10^8^ cells).

Finally, we tested whether AmvA could promote spermidine and spermine resistance in a bacterium lacking this MDE. *E. coli* BL21 cells overexpressing AmvA showed full growth in the presence of spermidine up to 20 mg/ml, while growth of the vector-only control strain decreased at ≥2.5 mg/ml spermidine (Fig 3E). AmvA also increased resistance to spermine to a lesser extent (Fig 3E). *AmvR is a spermine and spermidine-responsive repressor of amvA*

We next aimed to find out whether expression of *amvA* is regulated by its polyamine substrates, and if AmvR is responsible for this regulation. Previous work showed using qRT-PCR that *amvA* expression increases 5.7-fold in an *amvR* mutant (25). A reporter fusion vector, comprising the *amvR*-*amvA* intergenic region upstream of the GFP gene, showed significantly higher GFP expression in an *amvR* mutant of *A. baumannii* AB5075-UW (Fig 4A). This zeocin-resistant reporter construct showed slight toxicity and high baseline activity in *A. baumannii* AB5075-UW, so further polyamine induction experiments were conducted in *A. baumannii* ATCC17978 using a gentamicin-resistant reporter vector. Expression of *amvA*prom-GFP was not detected in late exponential phase, but increased slightly through stationary phase during a 4.5 hour incubation (Fig 4B). Putrescine and cadaverine at 10 mM did not affect expression of *amvA*prom-GFP, while 10 mM spermidine significantly increased GFP expression, with a 4-fold increase at the final time point. Spermine was toxic at 10 mM. No spermidine-dependent induction of *amvA* was observed in *A. baumannii* AB5075-UW *amvR* (Fig 4C).

**Figure 4.**
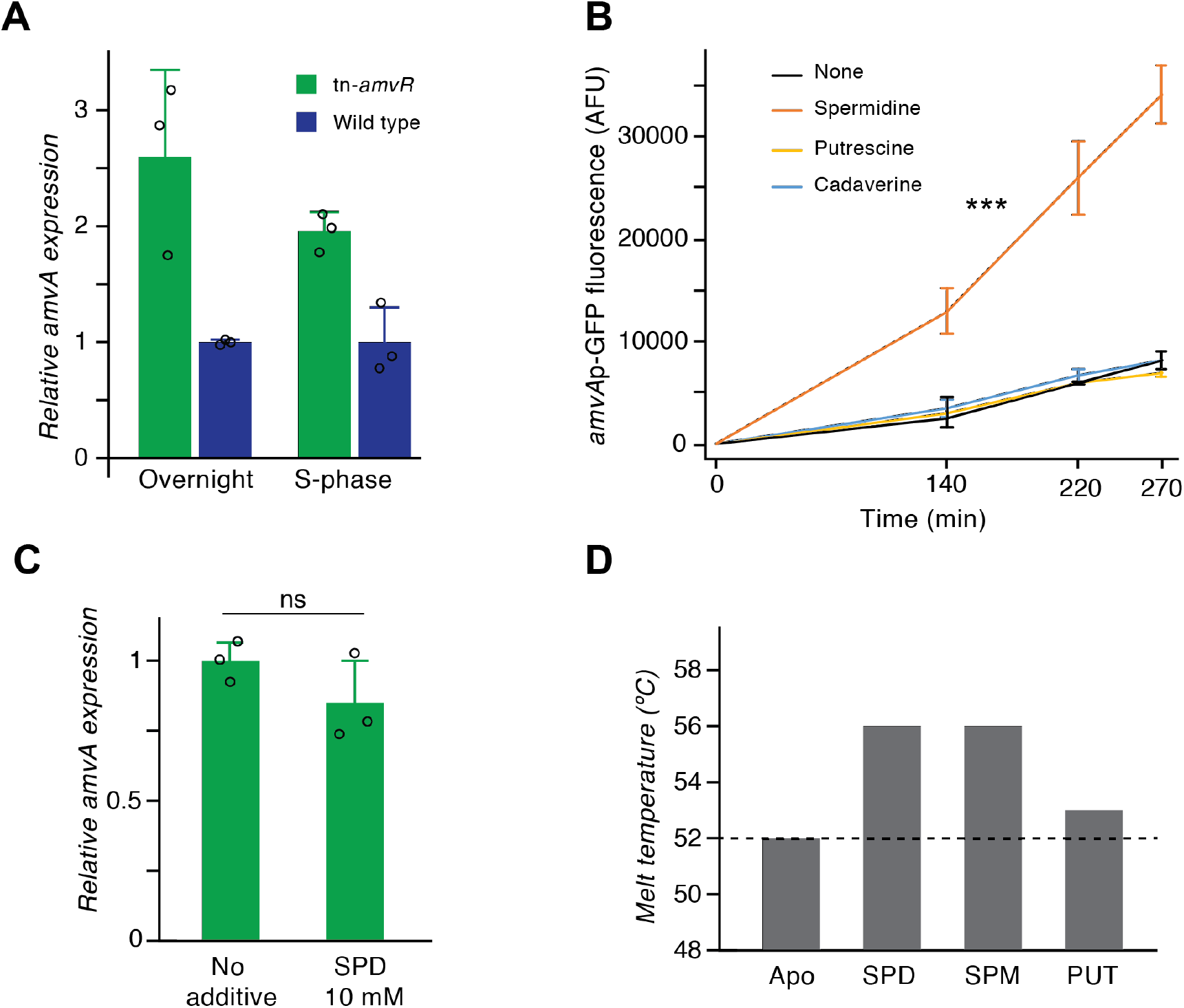
Regulation of *amvA* expression by AmvR and long-chain polyamines. (A) AmvR controls expression of *amvA*. Fluorescence of GFP in *A. baumannii* AB5075 or *A. baumannii* AB5075-tn-*amvR* carrying the *amvA* expression reporter vector pFLS45. The strain background (WT or tn-*amvR*) was identified as significant source of variation by two-way ANOVA, p< 0.0001. (B) Induction of *amvA* expression in *A. baumannii* ATCC17978 pFLS43 by 10mM CAD, PUT or SPD. Note that cells did not grow when 10 mM SPM was added. *** p < 0.0001, mixed repeated measures ANOVA (relative to WT). (C) Induction of *amvA*prom-GFP expression by SPD does not occur in AB5075 tn-*amvR*. (E) Shifts in melting temperature of purified AmvR in the presence of polyamines. Tm increases of 4 °C were seen in the presence of SPM and SPD, while PUT caused a 1°C shift. Experiments were performed in triplicate and gave identical Tm values.

We then tested whether a purified, recombinant form of AmvR has binding capacity for long-chain polyamines. Purified AmvR carrying a C-terminal hexahistidine tag was characterised by analytical SEC to be ~50kDa in solution, consistent with its native, stable dimeric form. Interactions of AmvR with polyamines in solution were monitored by perturbation of the protein’s melting temperature (Tm). Binding to a specific ligand will increase the thermal stability of a protein (27), and unfolding is observed as a dye response during differential scanning fluorimetry (DSF). As shown in Fig 4D, spermine and spermidine both elevated the thermal stability of AmvR by +4μC while putrescine caused little change (+1μC shift, i.e. below the significance threshold of the DSF experiment). These specific responses indicate that, in isolation, AmvR has capacity to form complexes with spermine or spermidine as ligands. Thus it appears that while AmvR represses *amvA* expression, this repression can be relieved by AmvR binding to long-chain polyamines.

## Discussion

Bacterial drug efflux pumps can have important physiological roles in addition to their resistance functions (5, 28). Here, we have investigated the effects of four of the predominant biological polyamines on *A. baumannii*, and identified a role for one of its major efflux pumps, AmvA, in the transport of long-chain polyamines. This is the first spermidine and spermine efflux system to be identified in *A. baumannii*.

Our findings support the hypothesis that polyamines have broad biological roles in *A. baumannii*. 499 genes had altered expression (representing 1/6 of the annotated CDS in AB5075), including many virulence-associated loci such as those encoding siderophores, pili and secretion systems (Fig 1, Fig 2, Table S3, S4). A limitation is that our transcriptomics was performed on cells shocked with high levels of exogenous polyamines, ranging from 18mM (spermine) to 56mM (putrescine). These concentrations were selected to show the full range of polyamine-responsive genes and to elicit strong induction of candidate transporters, but are higher than reported intracellular concentrations of polyamines in Gram-negative bacteria (eg. spermine ~6 mM, putrescine ~20 mM (29)), or in most host environments. The transcriptional responses defined here could arise from metabolic feed, biophysical modulation of DNA or RNA structure, activation of specific regulatory pathways, or a combination of all three. Experiments at physiological polyamine levels would help to define the biological roles of these molecules in more detail and determine whether *A. baumannii* utilises polyamines as specific virulence-regulating signals, as shown in some other bacterial species (17, 19).

Several lines of evidence indicate that long-chain polyamines are biological substrates of AmvA: 1) loss of this efflux pump increased spermidine accumulation and dramatically reduced spermine and spermidine tolerance, 2) *amvA* overexpression in *E. coli* increased spermidine and spermine resistance, 3) *amvA* expression is regulated in response to these molecules, and 4) regulation of *amvA* involves binding of these putative substrates to its cognate regulator, AmvR. Note that *amvA* expression is not induced by its other substrates including ciprofloxacin (30), deoxycholate (31), chlorhexidine (32) or benzalkonium chloride (33).

AmvA is clinically important in *A. baumannii*; it confers resistance to widely-used disinfectants (eg. chlorhexidine, benzalkonium), and in hospital-adapted strains its expression is increased (26). AmvA is a member of the DHA-2 (drug H^*^ antiporter-2) subfamily of MFS transporters. DHA-2 proteins comprise 14 transmembrane helices (34), and have a large, central hydrophobic cavity with several acidic residues that facilitate both proton and substrate binding (6, 35, 36), and determine substrate charge specificity (37). The substrate profile of AmvA includes a range of hydrophobic, cationic compounds with different degrees of protonation (26, 38). Spermine and spermidine are both long, flexible hydrophobic molecules with multiple protonated sites at physiological pH – it is likely that their transport requires particular active site properties, seen in DHA-2 proteins, that would also allow permit efflux of other substrates. AmvA is the first DHA-2 transporter shown to transport long-chain polyamines. Note that spermidine efflux has been demonstrated for the more distantly related DHA-1 transporter Blt, of *Bacillus subtilis* (21), which was also originally characterised for its drug-resistance activity (39).

We propose that the clinically-relevant anti-biocide activity of the well-known *A. baumannii* efflux pump AmvA stems from an ancestral role in homeostasis of long-chain polyamines. Interestingly, multiple bacterial pathogens including *Staphylococcus aureus*, *Proteus mirabilis* and members of the *Enterobacteriaceae* possess AmvA homologues (QacA, SmvA) which, like AmvA, confer increased biocide resistance (particularly to chlorhexidine) and are upregulated or have increased prevalence in clinical strains (40–43). It is tempting to speculate that clinically-important AmvA homologues may also have physiological roles in polyamine transport.

A further question is why *A. baumannii* would require specific efflux systems for spermine and spermidine? Such systems may protect from toxic levels of exogenous polyamines encountered in host environments. Polyamine levels can reach the mM range in plants, the gut, and in serum of patients suffering from some diseases (44–47). In particular, a recent study showed serum levels of spermidine and spermine at >500 µg/ml – close to their respective MICs in AB5075-UW *amvA* or *adeB* mutants (Fig 3B)(47). Alternatively, efflux of endogenous polyamines by AmvA (and perhaps AdeABC) may provide adaptive advantages unrelated to detoxification. For example, rapid putrescine efflux helps to counter osmotic stress in *E. coli* (48). *A. baumannii* produces a range of endogenous polyamines including spermidine (49–52), however their biological role(s) of are not yet fully understood.

The ability of *A. baumannii* to withstand biocides and antibiotics is a major barrier to the effective control of this pathogen. Here, we have shown that a key biocide resistance determinant, AmvA, is a spermidine and spermine efflux system, and provide preliminary evidence that AdeABC may share this activity. Together with previous findings on AceI (11), our work shows that three well-known *A. baumannii* efflux pumps, of three distinct transporter families, function in polyamine transport. All three of these pumps also transport the synthetic biocide chlorhexidine, which may be a common secondary substrate among polyamine efflux systems. Overall, our results suggest a strong link between homeostasis of the ancient, highly abundant polyamine class of molecules, and the broad and flexible efflux pump activity that allows bacterial pathogens – including *A. baumannii* – to survive treatment with disinfectants.

## Acknowledgements

We thank members of the Paulsen lab for useful discussions. This work was funded by a grant 1120298 from the National Health and Medical Research Council. KAH is supported by an Australian Research Council Future Fellowship FT180100123.

## Data availability

Raw RNA sequencing data associated with this study are available from the European Nucleotide Archive, accession PRJEB40527.

## Materials and methods

### Bacterial growth

Strains, plasmids and oligonucleotides used in this study are given in Table S1. *A. baumannii* was cultivated in Muller-Hinton (MH) broth at 37 °C for all experiments with the exception of reporter assays in *A. baumannii* AB5075-UW pFLS45, which were conducted in low-salt LB. When necessary, media were supplemented with polyamine salt compounds; spermidine tri-hydrochloride, putrescine di-hydrochloride, cadaverine di-hydrochloride or spermine tetra-hydrochloride. Antibiotics used were zeocin (150 µg/ml for *A. baumannii*; 25 µg/ml *E. coli*), gentamicin (10 µg/ml) and ampicillin (50 µg/ml).

### Antimicrobial susceptibility testing

MICs were determined by broth 2-fold serial dilution(53). Polyamine salts were added to the MH media (buffered to pH 7.8 using HEPES salt) to final concentrations of 80 mg/ml. Plates were incubated at 37 °C overnight and end-point growth at OD_600_ measured. *A. baumannii* strains were used in MIC experiments directly from overnight cultures, at an inoculum of ~10^6^ cells per ml, and plates were incubated without shaking. *E. coli* BL21 cells carrying either pTTQ18R6SH6 or pTTQ18R6SH6-AmvA (previously called AedF)(38) were prepared as follows: cells were grown overnight in MHII, subcultured at 1:10 and induced with 0.05mM IPTG for 1 hour at 37 °C 200rpm, and used to inoculate MIC broth dilution plates at approx. 2.5 × 10^5^ cfu/ml. Ampicillin and 0.05mM IPTG were included in media to maintain *amvA* expression, and plates were sealed with breathable film and incubated with shaking for 24 hours prior to OD_600_ measurement.

### RNA sequencing and analysis

*A. baumannii* AB5075-UW cells were grown overnight, and sub-cultured in MHII broth and grown to mid-exponential phase (5 ml, OD_600_ = 0.6). Cultures were treated with polyamine salts (1/8 MIC) for 30 min at 37 °C, with shaking. Control cultures contained no polyamine salt compounds. Cell pellets were recovered (5000g, 15 min at 4 ^0^C) and lysed in QIAzol reagent (Qiagen). Total RNA was isolated using the RNeasy mini kit (Qiagen) and DNA removed with DNaseI (TURBO DNA-free^TM^ kit, Invitrogen). rRNA removal and RNA sequencing were conducted by the Ramaciotti Centre (Sydney, Australia).

The raw data from the samples were analyzed using EDGE-PRO with Bowtie 2(54). Reads were normalized and differential expression analysis was conducted with the DESeq R package(55). A negative binomial model was used to test the significance of differential expression between control cells and polyamine-treated cells. Genes with adjusted p-values less than 0.05 and presenting at least 2-fold differences in expression were considered to be differentially expressed.

Functional enrichment analyses were conducted by first assigning COG (clusters of orthologous groups) and GO (gene ontology) terms to AB5075-UW genes using EggNOG mapper(56). Significantly enriched GO terms within the biological process, molecular function and cellular compartment ontologies were identified using the TopGO package with the ‘weight01’ algorithm and Fisher’s exact test.

### Serum resistance assays

Bacteria were grown overnight in MHII broth, subcultured 1:100 in fresh medium, and grown to late exponential phase (OD_600_ = 1). Cultures were then washed once in PBS and diluted 1:100 in sterile phosphate-buffered saline; 50 µl diluted culture was added to 100 µl pre-warmed human serum (Sigma) and incubated at 37 °C. Samples were taken at set time points, serially diluted and plated for enumeration of viable bacteria.

### Biofilm formation assays

2.5 ml cultures of *A. baumannii* AB5075-UW with or without polyamine supplementation were grown for 40 hours at 37 °C in 15 ml polystyrene tubes. Attached biofilm in the tubes was quantified by crystal violet staining as described(57).

### Pangenome analysis

All complete *A. baumannii* genome assemblies were downloaded from https://www.ncbi.nlm.nih.gov/assembly/ on March 26^th^, 2020. Genome sequences were annotated using prokka(58) version 1.14.6 with the following commands passed to the program: ‘prokka --prodigaltf prodigal_training.txt -- proteins AB5075_UWversion.gb --genus Acinetobacter --species baumannii’. The training model was generated using default settings, and the pipeline set to preferentially assign genes based on the *A. baumannii* AB5075-UW genome annotation. Gene content analysis across the population was conducted using Panaroo (59) version 1.1.2 with core gene alignment. Gene neighbourhood information was extracted from the final pangenome network using the get_neighbourhood script distributed with Panaroo.

### Spermidine transport assays

*A. baumannii* AB5075-UW and AB5075-UW tn-*amvA* were grown overnight in MHII broth, then diluted 1:100 in the same medium and grown to late exponential phase (OD_600_ = 1.0). Spermidine was added at 1/16 MIC to induce efflux pump expression, and cultures were incubated for a further 30 minutes. Cells were washed twice in assay buffer (20 mM HEPES pH 8.0, 145 mM NaCl, 5 mM KCl) and resuspended to a final OD_600_ of 1.0 in assay buffer supplemented with 0.5− (w/v) succinate to energise the cells. Accumulation assays were performed at room temperature with 1 ml resuspended cells. Bacterial samples were incubated with 10.8 pmol ^3^H-spermidine (46.1 mCi/mmol, Perkin-Elmer) for two minutes, then 200 µl of each reaction was immediately filtered through a 0.2 µM nitrocellulose membrane and washed with 2 ml assay buffer. Filters were placed in 5 ml PE Emulsifier-Safe scintillation fluid, and the amount of radiation retained on each filter (indicating intracellular ^3^H) was measured by liquid scintillation.

### DNA manipulations

To construct the reporter vector pFLS43 and pFLS45, the GFP marker from plasmid pDiGc was edited by overlap PCR using FS9 + FS10 (outer primers) and FS65 + FS66 (overlap primers) to remove its internal NdeI site, and introduced into pCR2.1 by TOPO cloning (Invitrogen). The *amvA*-*amvR* intergenic region from *A. baumannii* AB5075-UW was amplified using primers FS77 + FS78, and cloned upstream of the GFP-terminator fragment from pDiGc using the NdeI site. The promoter-GFP insert was subcloned from pCR2.1 into pVRL1 and pVRL1-Z using restriction enzymes SacI and XhoI. The AmvR expression vector was constructed by amplifying *amvR* from *A. baumannii* ATCC17978 genomic DNA using primers AmvR-F and AmvR-R, and cloning into the expression vector pTTQ_18RGSH6_.

### Promoter fusion experiments

Reporter plasmid pFLS45, comprising GFP under the control of the *amvA* promoter, was transformed into *A. baumannii* AB5075-UW and *A. baumannii* AB075 tn-*amvR* and maintained with zeocin selection in low-salt LB. Reporter plasmid pFLS43, which has the same structure but with gentamicin resistance, was transformed into *A. baumannii* ATCC17978 and maintained in gentamicin-supplemented MHII medium. Expression without induction (Fig 4A) was measured following growth in 10 ml cultures overnight or for six hours.

Induction by polyamines was measured by growing cells to late exponential phase in 10 ml cultures, then transferring cultures to a 96-well plate format (100 µl per well), adding polyamine salts, incubating the plate at 37 °C and measuring fluorescence and OD over time. For all experiments, expression was determined by measuring culture fluorescence (ex 485/ em 520) and OD_600_ in a 96-well plate in a Pherastar plate reader. GFP fluorescence/OD was calculated for each well, with cells lacking the reporter vector used as a background control. Results shown represent three biological replicates and six technical replicates.

### Expression and purification of AmvR

AmvR protein was expressed by growing BL21 pAmvR cells in ZYP-rich autoinduction medium (60) to an OD of 1.2-1.4 at 25 °C (approx. 24 hour incubation). Recovered cells were resuspended in 30 ml AmvR buffer (50 mM HEPES pH 7.5, 200 mM NaCl) supplemented with 5 mM imidazole and 5% glycerol and stored at −80C. Cell aliquots were thawed in the presence of lysozyme (1 µg/ml) and DNase I (5 µg/ml) and lysed by sonication. Clarified cell lysates (following 40 min centrifugation at 11,000 *x g* and 0.2 μm filtration) were loaded onto prepacked columns (1 ml) of Ni-Sepharose media (His trap, GE Healthcare) using a benchtop peristaltic pump. The column was then washed with 50 column volumes of AmvR buffer + 50 mM imidazole, and protein was eluted in AmvR buffer + 500mM imidazole.

Size exclusion chromatography was performed using a Superdex 200 HiLoad 16/60 column on an AKTA Pure liquid chromatography system, run at 1 ml/min. IMAC elution fractions were dialysed overnight into gel filtration buffer (50 mM HEPES pH 7.5, 300 mM NaCl, 10% glycerol) and then analysed by SEC in the same buffer. The molecular weight of AmvR in solution was estimated by comparing its elution volume to those of a set of molecular weight standards (HMW and LMW calibration sets, GE Healthcare).

### Differential scanning fluorimetry

DSF was performed according to standard methods (27). SYPRO Orange (100x, Invitrogen) was diluted into gel filtration buffer, and purified AmvR added to the dye mix. The final concentration of protein used in the screen was 1 mg/ml. 20 µl aliquots of AmvR-SYPRO orange mixture were transferred to a 96-well plate in triplicate, polyamine salts added at 0.2% (w/v), and the solutions gently mixed by plate centrifugation (1000 RPM, 1 min). The plate was heated at 1 °C/min over 25-95 °C in a real-time qPCR machine (Mx3005P, Strategene) and change in fluorescence intensity (dR, automatically baseline corrected) at 610 nm was measured. Derivative curves were used to calculate the transition midpoint and assign Tm. A change of ± 2 °C was seen to be significant for this work.

## Supplementary information

**Table S1.**
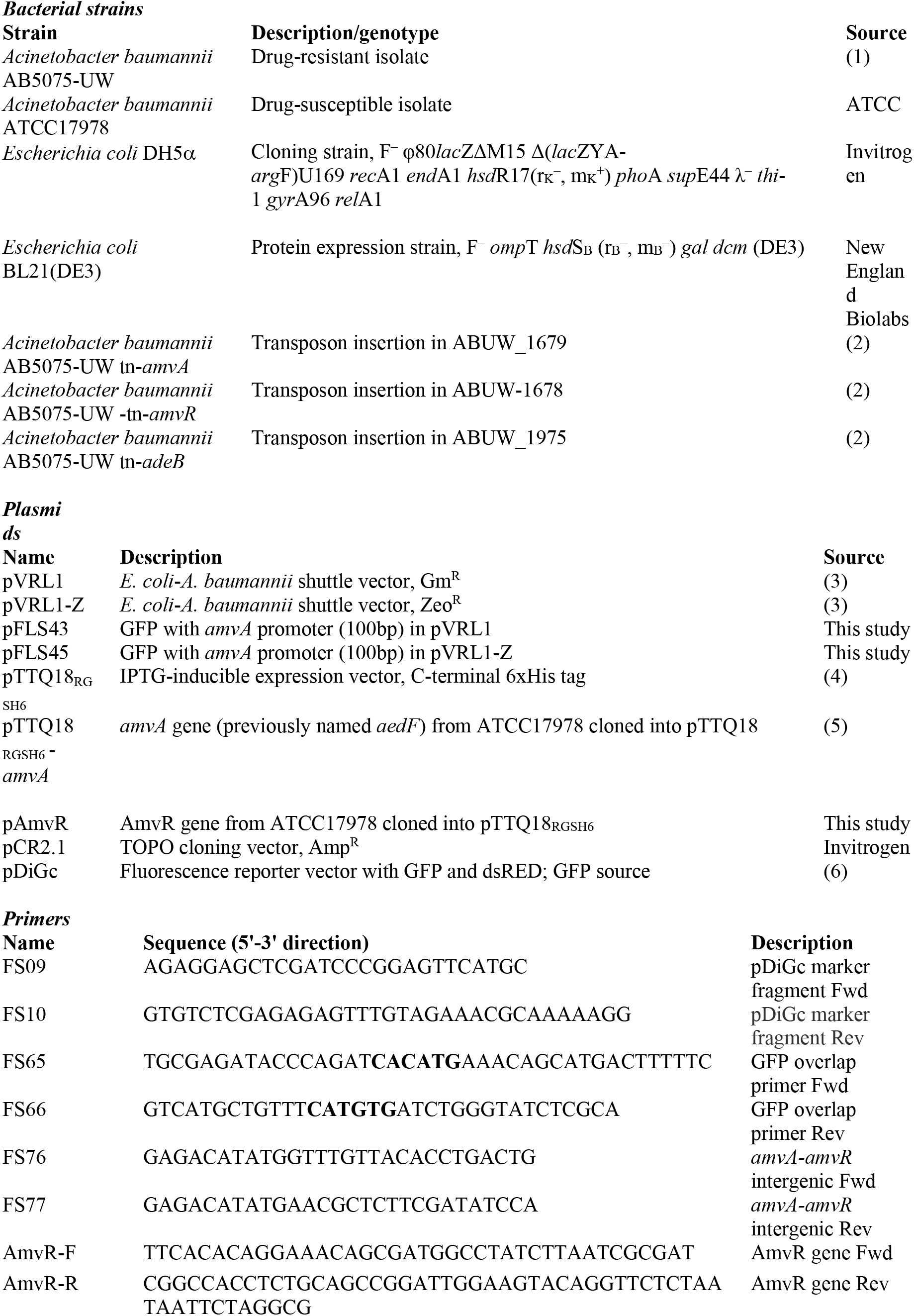
Strains, plasmids and oligonucleotides used in this study

**Table S2.** RNA sequence-reads and overall alignment rate in *ABUW-5075*

| Polyamines | Average sequence reads | Overall alignment rate |
| --- | --- | --- |
| Spermidine tri-hydrochloride | 11,090,209 ± 210,756 | 98.77% |
| Spermine tetra-hydrochloride | 10,414,484 ± 541,477 | 98.82% |
| Putrescine di-hydrochloride | 11,194,537 ± 1,155,240 | 99.19% |
| Cadaverine di-hydrochloride | 10,833,984 ± 487,424 | 98.96% |

**Table S3.** RNA-seq dataset (xlsx). Abbreviations: PUT = putrescine, CAD = cadaverine, SPD = spermidine, SPM = spermine. Spreadsheets containing full results, and polyamine-responsive genes.

**Table S4.** Polyamine-regulated genes of interest (xlsx). Abbreviations: PUT = putrescine, CAD = cadaverine, SPD = spermidine, SPM = spermine. Shown are locus tags, functional categories, specific regulation by polyamines, and any related enriched GO terms.

## References

1. O’Neill J. 2016. Tackling drug-resistant infections globally: Final report and recommendations. The Review on Antimicrobial Resistance.

2. Perry J, Waglechner N, Wright G. 2016. The prehistory of antibiotic resistance. Cold Spring Harb Perspect Med 6:1–9.

3. Davies J, Davies D. 2010. Origins and evolution of antibiotic resistance. Microbiol Mol Biol Rev 74:417–433.

4. Martinez JL, Sánchez MB, Martínez-Solano L, Hernandez A, Garmendia L, Fajardo A, Alvarez-Ortega C. 2009. Functional role of bacterial multidrug efflux pumps in microbial natural ecosystems. FEMS Microbiol Rev 33:430–449.

5. Blanco P, Hernando-Amado S, Reales-Calderon J, Corona F, Lira F, Alcalde-Rico M, Bernardini A, Sanchez M, Martinez J. 2016. Bacterial multidrug efflux pumps: Much more than antibiotic resistance determinants. Microorganisms 4:14.

6. Du D, Wang-Kan X, Neuberger A, van Veen HW, Pos KM, Piddock LJV, Luisi BF. 2018. Multidrug efflux pumps: structure, function and regulation. Nat Rev Microbiol 16:523–539.

7. Rice LB. 2008. Federal funding for the study of antimicrobial resistance in nosocomial pathogens: No ESKAPE. J Infect Dis 197:1079–1081.

8. Harding CM, Hennon SW, Feldman MF. 2017. Uncovering the mechanisms of *Acinetobacter baumannii* virulence. Nat Rev Microbiol 16:91.

9. Coyne S, Courvalin P, Périchon B. 2011. Efflux-mediated antibiotic resistance in *Acinetobacter* spp. Antimicrob Agents Chemother 55:947–953.

10. Peleg AY, Seifert H, Paterson DL. 2008. *Acinetobacter baumannii*: Emergence of a successful pathogen. Clin Microbiol Rev 21:538–582.

11. Hassan KA, Naidu V, Edgerton JR, Mettrick KA, Liu Q, Fahmy L, Li L, Jackson SM, Ahmad I, Sharples D, Henderson PJF, Paulsen IT. 2019. Short-chain diamines are the physiological substrates of PACE family efflux pumps. Proc Natl Acad Sci 116:18015–18020.

12. Michael AJ. 2016. Polyamines in eukaryotes, bacteria, and archaea. J Biol Chem 291:14896–14903.

13. Miller-Fleming L, Olin-Sandoval V, Campbell K, Ralser M. 2015. Remaining mysteries of molecular biology: the role of polyamines in the cell. J Mol Biol. Elsevier B.V.

14. Shah P, Swiatlo E. 2008. A multifaceted role for polyamines in bacterial pathogens. Mol Microbiol 68:4–16.

15. Ware D, Jiang Y, Lin W, Swiatlo E. 2006. Involvement of *potD* in *Streptococcus pneumoniae* polyamine transport and pathogenesis. Infect Immun 74:352–361.

16. Armbruster CE, Forsyth VS, Johnson AO, Smith SN, White AN, Brauer AL, Learman BS, Zhao L, Wu W, Anderson MT, Bachman MA, Mobley HLT. 2019. Twin arginine translocation, ammonia incorporation, and polyamine biosynthesis are crucial for *Proteus mirabilis* fitness during bloodstream infectionPLoS Pathogens.

17. Sobe RC, Bond WG, Wotanis CK, Zayner JP, Burriss MA, Fernandez N, Bruger EL, Waters CM, Neufeld HS, Karatan E. 2017. Spermine inhibits *Vibrio cholerae* biofilm formation through the NspS–MbaA polyamine signaling system. J Biol Chem 292:17025–17036.

18. Fang S Bin, Huang CJ, Huang CH, Wang KC, Chang NW, Pan HY, Fang HW, Huang M Te, Chen CK. 2017. *speG* is required for intracellular replication of *Salmonella* in various human cells and affects its polyamine metabolism and global transcriptomes. Front Microbiol 8:1–23.

19. Hobley L, Li B, Wood JL, Kim SH, Naidoo J, Ferreira AS, Khomutov M, Khomutov A, Stanley-Wall NR, Michael AJ. 2017. Spermidine promotes *Bacillus subtilis* biofilm formation by activating expression of the matrix regulator *slrR*. J Biol Chem 292:12041–12053.

20. Higashi K, Ishigure H, Demizu R, Uemura T, Nishino K, Yamaguchi A, Kashiwagi K, Igarashi K. 2008. Identification of a spermidine excretion protein complex (MdtJI) in *Escherichia coli.* J Bacteriol 190:872–878.

21. Woolridge DP, Vazquez-Laslop N, Markham PN, Chevalier MS, Gerner EW, Neyfakh AA. 1997. Efflux of the natural polyamine spermidine facilitated by the *Bacillus subtilis* multidrug transporter Blt. J Biol Chem 272:8864–8866.

22. Alexa A. 2013. Gene Set Enrichment Analysis. Encycl Syst Biol 806–806.

23. Morris FC, Dexter C, Kostoulias X, Uddin MI, Peleg AY. 2019. The mechanisms of disease caused by *Acinetobacter baumannii.* Front Microbiol 10.

24. Fouts DE. 2006. Phage_Finder: Automated identification and classification of prophage regions in complete bacterial genome sequences. Nucleic Acids Res 34:5839–5851.

25. Hassan KA, Cain AK, Huang T, Liu Q, Elbourne LDH, Boinett CJ, Brzoska AJ, Li L, Ostrowski M, Nhu NTK, Nhu TDH, Baker S, Parkhill J, Paulsen IT. 2016. Fluorescence-based flow sorting in parallel with transposon insertion site sequencing identifies multidrug efflux systems in *Acinetobacter baumannii*. MBio 7:3–8.

26. Rajamohan G, Srinivasan VB, Gebreyes WA. 2010. Molecular and functional characterization of a novel efflux pump, AmvA, mediating antimicrobial and disinfectant resistance in *Acinetobacter baumannii.* J Antimicrob Chemother 65:1919–1925.

27. Niesen FH, Berglund H, Vedadi M. 2007. The use of differential scanning fluorimetry to detect ligand interactions that promote protein stability. Nat Protoc 2:2212–2221.

28. Alcalde-Rico M, Hernando-Amado S, Blanco P, Martinez JL. 2016. Multidrug efflux pumps at the crossroad between antibiotic resistance and bacterial virulence. Front Microbiol 7:1–14.

29. Tabor CW, Tabor H. 1985. Polyamines in microorganisms. Microbiol Rev 49:81–99.

30. Geisinger E, Vargas-Cuebas G, Mortman NJ, Syal S, Dai Y, Wainwright EL, Lazinski D, Wood S, Zhu Z, Anthony J, van Opijnen T, Isberg RR. 2019. The landscape of phenotypic and transcriptional responses to ciprofloxacin in *Acinetobacter baumannii:* Acquired resistance alleles modulate drug-induced SOS response and prophage replication. MBio 10:1–19.

31. López M, Blasco L, Gato E, Perez A, Fernández-Garcia L, Martínez-Martinez L, Fernández-Cuenca F, Rodríguez-Baño J, Pascual A, Bou G, Tomás M, GEIH-GEMARA (SEIMC). 2017. Response to bile salts in clinical strains of *Acinetobacter baumannii* lacking the *adeABC* efflux pump: Virulence associated with quorum sensing. Front Cell Infect Microbiol 7:1–13.

32. Hassan KA, Jackson SM, Penesyan A, Patching SG, Tetu SG, Eijkelkamp BA, Brown MH, Henderson PJF, Paulsen IT. 2013. Transcriptomic and biochemical analyses identify a family of chlorhexidine efflux proteins. Proc Natl Acad Sci 110:20254–20259.

33. Knauf GA, Cunningham AL, Kazi MI, Riddington IM, Crofts AA, Cattoir V, Trent MS, Davies BW. 2018. Exploring the antimicrobial action of quaternary amines against *Acinetobacter baumannii.* MBio 9:1–13.

34. Reddy VS, Shlykov MA, Castillo R, Sun EI, Saier MH. 2012. The major facilitator superfamily (MFS) revisited. FEBS J 279:2022–2035.

35. Paulsen IT. 2003. Multidrug efflux pumps and resistance: Regulation and evolution. Curr Opin Microbiol 6:446–451.

36. Majumder P, Khare S, Athreya A, Hussain N, Gulati A, Penmatsa A. 2019. Dissection of protonation sites for antibacterial recognition and transport in QacA, a multi-drug efflux transporter. J Mol Biol 431:2163–2179.

37. Paulsen IT, Brown MH, Littlejohn TG, Mitchell BA, Skurray RA. 1996. Multidrug resistance proteins QacA and QacB from *Staphylococcus aureus*: Membrane topology and identification of residues involved in substrate specificity. Proc Natl Acad Sci U S A 93:3630–3635.

38. Hassan KA, Brzoska AJ, Wilson NL, Eijkelkamp BA, Brown MH, Paulsen IT. 2011. Roles of DHA2 family transporters in drug resistance and iron homeostasis in *Acinetobacter* spp. J Mol Microbiol Biotechnol 20:116–124.

39. Ahmed M, Lyass L, Markham PN, Taylor SS, Vazquez-Laslop N, Neyfakh AA. 1995. Two highly similar multidrug transporters of *Bacillus subtilis* whose expression is differentially regulated. J Bacteriol 177:3904–3910.

40. Wand ME, Jamshidi S, Bock LJ, Rahman KM, Sutton JM. 2019. SmvA is an important efflux pump for cationic biocides in *Klebsiella pneumoniae* and other Enterobacteriaceae. Sci Rep 9:1–11.

41. Pelling H, Bock LJ, Nzakizwanayo J, Wand ME, Denham EL, MacFarlane WM, Sutton JM, Jones B V. 2019. Derepression of the *smvA* efflux system arises in clinical isolates of *Proteus mirabilis* and reduces susceptibility to chlorhexidine and other biocides. Antimicrob Agents Chemother 63:1–15.

42. Villagra NA, Hidalgo AA, Santiviago CA, Saavedra CP, Mora GC. 2008. SmvA, and not AcrB, is the major efflux pump for acriflavine and related compounds in *Salmonella enterica* serovar Typhimurium. J Antimicrob Chemother 62:1273–1276.

43. Grkovic S, Brown MH, Roberts NJ, Paulsen IT, Skurray RA. 1998. QacR is a repressor protein that regulates expression of the *Staphylococcus aureus* multidrug efflux pump QacA. J Biol Chem 273:18665–18673.

44. Bartos F, Bartos D, Grettie DP, Campbell RA. 1977. Polyamine levels in normal human serum. Biochem Biophys Res Commun 75:1689–1699.

45. Kingsnorth AN, Lumsden AB, Wallace HM. 1984. Polyamines in colorectal cancer. Br J Surg 71:791–794.

46. Ramos-Molina B, Queipo-Ortuño MI, Lambertos A, Tinahones FJ, Peñafiel R. 2019. Dietary and gut microbiota polyamines in obesity- and age-related diseases. Front Nutr 6:1–15.

47. Ibrahim SA, Zainulabdeen JA, Jasim HM. 2018. The significance of spermidine and spermine in association with atherosclerosis in sera of Iraqi patients. Biomed Pharmacol J 11:1389–1396.

48. Schiller D, Kruse D, Kneifel H, Kramer R, Burkovski A. 2000. Polyamine transport and role of *potE* in response to osmotic stress in *Escherichia coli*. J Bacteriol 182:6247–6249.

49. Hamana K, Matsuzaki S. 1992. Diaminopropane occurs ubiquitously in *Acinetobacter* as the major polyamine. J Gen Appl Microbiol 38:191–194.

50. Skiebe E, de Berardinis V, Morczinek P, Kerrinnes T, Faber F, Lepka D, Hammer B, Zimmermann O, Ziesing S, Wichelhaus TA, Hunfeld KP, Borgmann S, Gröbner S, Higgins PG, Seifert H, Busse HJ, Witte W, Pfeifer Y, Wilharm G. 2012. Surface-associated motility, a common trait of clinical isolates of *Acinetobacter baumannii*, depends on 1,3-diaminopropane. Int J Med Microbiol 302:117–128.

51. Auling G, Pilz F, Busse HJ, Karrasch S, Streichan M, Schon G. 1991. Analysis of the polyphosphate-accumulating microflora in phosphorus-eliminating, anaerobic-aerobic activated sludge systems by using diaminopropane as a biomarker for rapid estimation of *Acinetobacter* spp. Appl Environ Microbiol 57:3585–3592.

52. Kämpfer P, Bark K, Busse HJ, Auling G, Dott W. 1992. Numerical and chemotaxonomy of polyphosphate accumulating *Acinetobacter* strains with high polyphosphate: AMP phosphotransferase (PPAT) Aativity. Syst Appl Microbiol 15:409–419.

53. Wiegand I, Hilpert K, Hancock REW. 2008. Agar and broth dilution methods to determine the minimal inhibitory concentration (MIC) of antimicrobial substances. Nat Protoc 3:163–175.

54. Magoc T, Wood D, Salzberg SL. 2013. EDGE-pro: Estimated Degree of Gene Expression in Prokaryotic Genomes. Evol Bioinform Online. 9:127–136.

55. Love MI, Huber W, Anders S. 2014. Moderated estimation of fold change and dispersion for RNA-seq data with DESeq2. Genome Biol 15:550.

56. Huerta-Cepas J, Forslund K, Coelho LP, Szklarczyk D, Jensen LJ, Von Mering C, Bork P. 2017. Fast genome-wide functional annotation through orthology assignment by eggNOG-mapper. Mol Biol Evol 34:2115–2122.

57. Biswas I, Mettlach J. 2013. A simple static biofilm assay for *Acinetobacter baumannii*, p. 159–166. In Biswas, I, Rather, PN (eds.), Acinetobacter baumannii Methods and Protocols.

58. Seemann T. 2014. Prokka: rapid prokaryotic genome annotation. Bioinformatics 30:2068–2069.

59. Tonkin-Hill G, MacAlasdair N, Ruis C, Weimann A, Horesh G, Lees JA, Gladstone RA, Lo SW, Beaudoin C, Floto RA, Frost SDW, Corander J, Bentley SD, Parkhill J. 2020. Producing Polished Prokaryotic Pangenomes with the Panaroo Pipeline. Genome Biol 21:180

60. Studier FW. 2005. Protein production by auto-induction in high-density shaking cultures. Protein Expr Purif 41:207–234.

## Supplementary references

1. Jacobs AC, Thompson MG, Black CC, Kessler JL, Clark LP, McQueary CN, Gancz HY, Corey BW, Moon JK, Si Y, Owen MT, Hallock JD, Kwak YI, Summers A, Li CZ, Rasko DA, Penwell WF, Honnold CL, Wise MC, Waterman PE, Lesho EP, Stewart RL, Actis LA, Palys TJ, Craft DW, Zurawski D V. 2014. AB5075, a Highly Virulent Isolate of Acinetobacter baumannii, as a Model Strain for the Evaluation of Pathogenesis and Antimicrobial Treatments. MBio 5:e01076–14.

2. Gallagher LA, Ramage E, Weiss EJ, Radey M, Hayden HS, Held KG, Huse HK, Zurawski D V., Brittnacher MJ, Manoil C. 2015. Resources for genetic and genomic analysis of emerging pathogen Acinetobacter baumannii. J Bacteriol 197:2027–2035.

3. Lucidi M, Runci F, Rampioni G, Frangipani E, Leoni L, Visca P. 2018. New shuttle vectors for gene cloning and expression in multidrug-resistant Acinetobacter species. Antimicrob Agents Chemother 62:1–19.

4. Stark MJ. 1987. Multicopy expression vectors carrying the lac repressor gene for regulated high-level expression of genes in Escherichia coli. Gene 51:255–267.

5. Hassan KA, Brzoska AJ, Wilson NL, Eijkelkamp BA, Brown MH, Paulsen IT. 2011. Roles of DHA2 family transporters in drug resistance and iron homeostasis in Acinetobacter spp. J Mol Microbiol Biotechnol 20:116–124.

6. Helaine S, Thompson JA, Watson KG, Liu M, Boyle C, Holden DW. 2010. Dynamics of intracellular bacterial replication at the single cell level. Proc Natl Acad Sci U S A 107:3746–3751.

